# Successful Correction of ALD Patient-derived iPSCs Using CRISPR/Cas9

**DOI:** 10.1101/2020.02.23.962118

**Authors:** Eul Sik Jung, Zhejiu Quan, Mi-Yoon Chang, Wonjun Hong, Ji Hun Kim, Seung Hyun Kim, Seungkwon You, Dae-Sung Kim, Jiho Jang, Sang-Hun Lee, Hyongbum (Henry) Kim, Hoon Chul Kang

**Author notes:** Corresponding author: Hoon Chul, Kang, MD, PhD. These authors contributed equally to this work (Eul Sik Jung, Zhejiu Quan, Mi-Yoon Chang).

## Abstract

X-linked adrenoleukodystrophy (ALD) caused by the *ABCD1* mutation, is the most common inherited peroxisomal disease. It is characterized by three phenotypes: inflammatory cerebral demyelination, progressive myelopathy, and adrenal insufficiency, but there is no genotype-phenotype correlation. Hematopoietic stem cell transplantation can only be used in a few patients in the early phase of cerebral inflammation; therefore, most affected patients have no curative option. Previously, we reported the generation of an ALD patient-derived iPSC model and its differentiation to oligodendrocytes. In this study, we have performed the first genome editing of ALD patient-derived iPSCs using homology-directed repair (HDR). The mutation site, c.1534G>A [GenBank: NM_000033.4], was corrected by introducing ssODN and the CRISPR/Cas9 system. The cell line exhibited normal ALD protein expression following genome editing. We differentiated the intermediate oligodendrocytes from mutation-corrected iPSCs and the metabolic derangement of ALD tended to correct but was not statistically significant. Mutation-corrected iPSCs from ALD patient can be used in research into the pathophysiology of and therapeutics for ALD.

## Introduction

X-linked adrenoleukodystrophy (ALD, OMIM:300100) is a rare genetic disease with an overall frequency of 1:17,000 [1]. The clinical manifestation of ALD is expressed as inflammatory cerebral demyelination, progressive myelopathy, and endocrine dysfunction such as an adrenal insufficiency or gonadal dysfunction [2]. Very long chain fatty acids (VLCFAs) are accumulated in the cytoplasm due to a defect in the ALD protein, encoded by the *ABCD1* gene. However, there is no correlation between clinical manifestation and the amount of VLCFA, or even the genotype [3]. For example, even identical twins presented a different phenotype [4]. The only common feature in ALD patients is the *ABCD1* mutation. The inflammatory cerebral demyelination seen in ALD patients deteriorates rapidly, and the patient dies or enters a vegetative state. The only way to halt demyelination is via hematopoietic stem cell transplantation (HSCT), but only a few patients at the early stage of demyelination are eligible to receive HSCT [5].

Induced pluripotent stem cells (iPSCs) were introduced in 2006 and have since been applied in research and therapeutic purposes [6]. Through the introduction of reprogramming factors, somatic cells can be altered into embryonic stem cells (ESCs)-like cells, with morphology, gene expression profile, and pluripotency characteristic of ESCs. Personalized iPSC can be generated by reprogramming patient somatic cells and can differentiate into a wide variety of other cell types, including neural stem cells. In case of rare diseases that lack *in vitro* or *in vivo* experimental models, patient-derived iPSCs enable researches to study pathophysiology, conduct high-throughput drug screening, and develop stem cell therapeutic strategies. Recently, clinical trials have been performed using iPSCs as therapeutics, although there are some limitations [7]. We reported the development of an ALD iPSC model, and observed significant differences between normal and oligodendrocytes originated from ALD-derived iPSCs [8]. The VLCFA metabolic ratio and expression level of *ABCD2* was relatively higher, but expression of *ELOVL1* was lower, in the ALD group.

CRISPR/Cas9 (clustered regularly interspaced short palindromic repeats and CRISPR-associated protein 9) can bind to a specific DNA region, and/or make a double-strand break or a nick. It can generate an *indel,* leading to gene disruption, or enable homology-directed repair (HDR), leading to a gene insertion or precise correction [9]. Therefore, many attempts have been made to generate a disease model or to design therapeutic materials using CRISPR/Cas9-mediated iPSC genome editing [10]. In particular, a corrected patient-derived iPSC using genome editing technology has no risk of immune rejection upon transplantation [10].

Here, we have generated an ALD patient-derived iPSC and performed successful correction of the disease-causing mutation through HDR, by introducing the ssODN and the CRISPR/Cas9 system. We made intermediate oligodendrocytes from mutation-corrected iPSc. It was confirmed by Sanger sequencing and reinstated normal *ABCD1* transcription. There were minimal and insignificant off-targets in all mutation-corrected iPSC lines. VLCFA accumulation in oligodendrocytes from mutation corrected iPSCs tended to decrease in comparison to ALD oligodendrocytes, but it was statistically insignificant.

## Materials and Methods

### Ethics statement

All experiments were conducted under the supervision of the Human Research Protection Center, Yonsei University College of Medicine, and followed the guidelines of the Institutional Review Board (Approval No. 4-2016-0194). The patient’s skin sample was donated voluntarily.

### Establishment of iPSC lines from human fibroblasts

Human fibroblast cell lines were isolated from a patient carrying the *ABCD1* missense mutation c.1534G>A [GenBank: NM_000033.4] by skin biopsy. The patient is 34 years old and has myelopathy without cerebral demyelination, called adrenomyeloneuropathy (AMN). For generating iPSCs, patient-derived fibroblasts were infected with Sendai virus using the CytoTune-iPS 2.0 Sendai Reprogramming Kit (Invitrogen, CA, US) according to the manufacturer’s instruction, and fibroblasts unaffected by ALD were used as control (designated WT) fibroblasts (ATCC, VA, US; Cat. No.: CRL-2522). Briefly, fibroblasts were plated with low-glucose Dulbecco modified Eagle medium (DMEM) (Thermo Fisher Scientific, MA, US) on the day of transduction. We transduced cells using the CytoTune-iPS 2.0 Sendai Reprogramming vector according to the manufacturer’s instructions, and then incubated overnight. The culture medium was replaced every day before transfer. After 30 days, iPSC colonies were transferred to mouse embryonic fibroblast feeder cells (ATCC, VA, US; Cat. No.: CRL-1503) for expansion and maintained with conventional human embryonic stem cell medium^7^. To assess pluripotency, established iPSC lines were characterized by alkaline phosphatase staining (Vector Laboratories, CA, US; Cat. No. SK-5300) and immunocytochemistry, including staining for SSEA4 (Merck Millipore, Germany; Cat. No. MAB4304), OCT4 (Merck Millipore, Germany; Cat. No. AB3209), SOX2 (Merck Millipore, Germany; Cat. No. AB5603), NANOG (Abcam, Canada; Cat. No. AB80892), TRA-1-60 (Merck Millipore, Germany; Cat. No. MAB4360), and TRA-1-81 (Merck Millipore, Germany; Cat. No. MAB4381).

### Plasmids containing Cas9, sgRNA, and ssODN

The vector, including SpCas9 and selection cassettes of Red Florescent Protein (RFP) and the puromycin resistance gene, was purchased (pCas-Puro^R^/RFP; Toolgen, South Korea). pRGEN (Toolgen, South Korea) containing sgRNA under the U6 promotor was used for cloning pRGEN_ABCD1 (sgRNA sequences, AGCTCCCTGTTCCGGATCC). The singlestranded oligodeoxynucleotides (ssODNs) included five base pairs that were different from the patient’s genomic sequences (Supplementary sequences): three of these formed a silent mutation site to prevent re-cleavage after homology-directed repair (HDR), another was part of an artificial silent restriction site of Apa1, and the last one was for the correction of disease-causing missense mutation. ssODNs were synthesized by Integrated DNA Technology (IA, US).

### Genome editing using HDR in patient-derived iPSCs

We used feeder-free conditions for the genome editing experiment. Essential 8 medium (Life Technologies, CA, US) was used for feeder-free culture. Transfections were performed using the NEPA21 electroporator (Nepagene, Japan) and cells were then plated onto a Matrigel-coated 60-mm dish with Essential 8 medium. After allowing the cells to settle for 3 h, single isolated cells were marked with 3-mm circle markers on the bottom of the plate. Only colonies grown within the marked circles were used for further analysis. RFP+ cells appeared within 24 h after electroporation. 12h after transfection, cells were subjected to puromycin selection (1 μg/ml) for 3 h and the medium was subsequently changed to remove puromycin. Colonies after puromycin selection were maintained for 10 days. Then, growing colonies were manually picked, triturated, and plated onto 24-well plates. The triturated colonies were cultured for 4 more days until sufficient confluency. From all colonies in each well, we collected only half portion of each colony by Pasteur pipette and mixed the collected half colonies into one tube. Then, the genomic DNA from the mixture was extracted and analyzed using Apa1 digestion and Sanger sequencing (described below). We selected colonies that were positive for gene correction, and transferred these to 24-well plates. At this step, one single colony was plated in each well. We repeated this procedure to get pure mutation-corrected iPSCs.

### Differentiation of oligodendrocytes from human iPSCs

Human iPSCs were dissociated to single cells and seeded on Matrigel coated 6 well plate on day −2. On day 0, medium was switched to N2B27 medium (DMEM/F12 (Hyclone, UT, US; Cat. No. #SH30023.01) supplemented with N2 (Gibco; Cat. No. #17502048) (1:100), B27(Gibco; Cat. No. #12587010) (1:50), glutamax (1:100), NEAA (1:100), betamercaptoethanol (1:1000) and Pen/Strep (Gibco) (1:100)) supplemented with 10μM SB431542 (Tocris, UK; Cat. No. #1614), 1 μM LDN193189, 25μg/ml human insulin (Sigma-Aldrich, MO, US; Cat. No. #I9278) and 100 nM RA (Sigma-Aldrich, MO, US; Cat. No. #R2625) until day 8, when SB431542 and LDN193189 (Tocris, UK; Cat. No. #6053) were withdrawn and smoothened agonist 1μM SAG (Cayman Chemical, MI, US; Cat. No #11914) added. On day 12, cells were dissociated, plated, and transduced with viral vectors. Next day, medium was changed to OL differentiation medium (ODM) with the addition of 1 mg/mL doxycycline, that supplemented, IGF1 (Peprotech, NJ, US; Cat. No. #100-11) 10ng / ml, HGF (Peprotech, NJ, US; Cat. No. #100-39H) 5ng / ml, cAMP (Sigma-Aldrich, MO, US; Cat. No. #A6885) 1uM, NT-3 10 ng/ml, T3 (Sigma-Aldrich, MO, US; Cat. No. #T6397) 60ng / ml, PDGFaa (Peprotech NJ, US; Cat. No. #100-13A) 10ng / Consists of ml, Biotin (Sigma-Aldrich, MO, US; Cat. No. #B4501) 100 ng/ml. Cells were maintained for 10-12 days.

For oligodendrocyte (OL) differentiation, SOX10 lentivirus was used. We selected mammalian transcription factors (TFs) involved in OL differentiation: SOX10 [11]. The lentiviral vector Teto-O-FUW-Sox10 (Addgene, MA, USA; Cat. No. 45843) and FUW-M2-rtTA (Addgene, MA, US; Cat. No. 20342) were co-transfected with the envelope plasmids pRSV-Rev (Addgene, MA, US; Cat. No. 12253), pMDLg/pRRE (Addgene, MA, US; Cat. No. 12251) and pMD2. G (Addgene, MA, US; Cat No.: 12259) into the 293FT cell line (Invitrogen, CA, US; Cat. No. R70007) using lipofectamine 3000 transfection reagent (Invitrogen, CA, US; Cat. No.: L3000-001). Supernatants containing the lentiviral particles were collected after 48hr, filtered through a 0.45μm filter (Merck Millipore, Germany) and stored at −80 °C for future use.

In the case of immunocytochemistry, fix cells with 4% PFA on ODM d-10 and proceed with blocking with 5% donkey serum. In the case of O4 antibody (Merck Millipore, Germany; Cat. No. MAB1326), dilution was performed at 1:200, followed by incubation at 4°C during overnight, followed by 30 min RT incubation with anti-Mouse IgM 594 (Invitrogen, CA, US; Cat. No. A-21044) 1: 500. In the case of MBP antibody (Merck Millipore, Germany; Cat. No. MAB 386), after dilution at 1:50 with 0.2% Triton-X 100 for 20 min permeabilization, and incubation at 4°C overnight, anti-Rat IgG (Invitrogen, CA, US; Cat. No. A-21208) 1:500 for 30 min RT Proceed with incubation. In both cases, DAPI stained and observed under a fluorescence microscope.

### Genomic DNA extraction and PCR

Genomic DNA (gDNA) was isolated by using DNeasy Blood & Tissue Kit (QIAGEN, Germany; Cat. No. 69504) according to the manufacturer’s protocol. 50-100ng of gDNA used for each PCR reaction with pfu Taq (Bioneer, Korea; Cat. No. K-2303). 742bp around exon 6 of *ABCD1* gene was by amplified PCR (Forward primer; CTGTGGCAGAATAGGCCCTT, Reverse primer; CTCCCCCAAGATACTCTGCG). Each amplified PCR samples were purified with agarose gel and analyzed by Sanger sequencing.

### Western blot

Cell lysates were prepared using RIPA buffer (Rockland Immunochemicals, PA, US; Cat. No. MB-030-0050). Samples were centrifuged at 12,000 × g for 10 min at 4°C and the supernatant was isolated and quantified using a Pierce BCA protein assay kit (Thermo Fisher Scientific, MA, US). Loading buffer (4×) was added into the samples, and they were boiled for 5 min at 90°C. The extracted protein samples were separated on an 10% SDS-PAGE gel by electrophoresis and transferred to polyvinylidene fluoride membranes. Membranes were blocked in TBS-T (1% Tween 20) with 5% BSA for 30 min and incubated with primary antibody against human ABCD1 (1:1,000; Origene, MD, US; Cat. No. TA803208) and β-actin (1:1,000; Santacruz, TX, US; Cat. No. sc-47778) for 40 h at 4°C. After washing, membranes were incubated with anti-mouse antibody (1:5,000; Merck Millipore, Germany; Cat. No. AP124P). After incubation, blots were visualized using the ECL western blotting detection reagent (GE Healthcare, IL, US; Cat. No. RPN2106) on an electrochemiluminescence^10^ instrument (Amersham Biosciences, UK; Cat. No. RPN210).

### Off-target analysis

At first, we performed *in silico* off-target analysis using Cas-OFFinder (http://www.rgenome.net/cas-offinder/)[12]. Off-target analysis permitted two base pair mismatches and one DNA bulge in Cas-OFFinder. All possible on-target and off-target mutations were also analyzed by whole exome sequencing (WES). The exons were captured using the Agilent Sureselect kit, version V6, which covers the exomes of more than 20,000 genes. These were sequenced on an Illumina Novaseq 6000 sequencing system and the reads were mapped to the human reference genome GRCh37. We called variants using the Haplotype Caller algorithm from the Genome Analysis Toolkit (GATK) (3.8-0) (http://www.broadinstitute.org/gatk/).

### Lipid analysis

For VLCFA analysis, cells were accumulated with accutase on OL differentiation medium (ODM) day 10, and MACs sorted using Anti-O4 MicroBeads (Miltenyi Biotec, Germany) and LS columns (Miltenyi Biotec, Germany). In the case of magnetic activated cells (MACs) sorting, the experiment was conducted as recommended by manufacture. Briefly, 60ul of MACs buffer and 20 μl of Anti-O4 MicroBeads per 1 x 10^7^ cells were incubated for 15 minutes on ice, and after washing, LS column was mounted on QuadroMACS ™ Separators and sorted by washing step. After sorting, 2.0×10^5^ cells were mixed with 450 μl of PBS and analyzed by VLCFA through GC-MS in SCL.

VLCFA analysis was performed at Seoul Clinical Laboratories, Seoul Medical Science Institute, Korea (http://www.scllab.co.kr). VFCFA analysis was performed using gas chromatography/mass spectrometry (GC/MS), as previously noted[8]. For VLCFA analysis, harvested cells were sonicated and mixed with 25% methylene chloride in methanol. Next, 200 ul of acetyl chloride was added, and the mixture was incubated at 75 °C for 1 h. After cooling, 4 ml of 7% K_2_CO_3_ was added to stop the reaction by neutralization. Subsequently, 5 ml of hexane was added and vortexed, and the mixture was then centrifuged. The hexane layer was removed to a new glass tube and mixed with 2 ml of acetonitrile. The isolated layer was moved to a new glass tube, then dried under a gentle nitrogen stream at 40°C. Then, 50 μl of hexane was added to dry the residue, which was analyzed using a gas chromatography/mass spectrometry detector (GC/MSD; Agilent Technologies, US).

For lysophosphatidyl choline (LPC) analysis, 150 μl of methanol was added to 50 μl of cell pellets with PBS. The mixture was incubated for 30 minutes at room temperature and centrifuged. The separated layer was transferred to a liquid chromatography vial and analyzed. LPC was analyzed by electrospray ionization (ESI) with tandem mass spectrometry, using an HPLC system (Agilent 1260, Agilent, US) and MS/MS system (5500 QTRAP, AB SCIEX, US).

## Results

### Generation of patient-derived iPSCs

Human fibroblasts were isolated from a biopsied patient’s skin who has the *ABCD1* mutation c.1534G>A [GenBank: NM_000033.3] (designated DS). DS fibroblasts were transduced with Sendai viral vectors expressing the reprogramming factors Oct, Sox2, Klf4, and c-Myc, along with WT fibroblasts. Transduced cells were plated on mouse feeder cell culture dish. After 20 to 30 days, emerging iPSC colonies were separately transferred to new fresh mouse feeder cells, then expanded for 6 days. The pluripotency of the established iPSC lines was verified by alkaline phosphatase and immunocytochemistry assays (Fig 1A). Intense alkaline phosphatase activity was observed. Pluripotency markers, including OCT3/4, SOX2, TRA-1-81, SSEA4, TRA-1-60, and NANOG, were expressed. The karyotype was normal without abnormalities in the number or structure of chromosomes. Sanger sequencing for genomic DNA confirmed the c.1534G>A mutation (Fig 1B).

**Fig 1.**
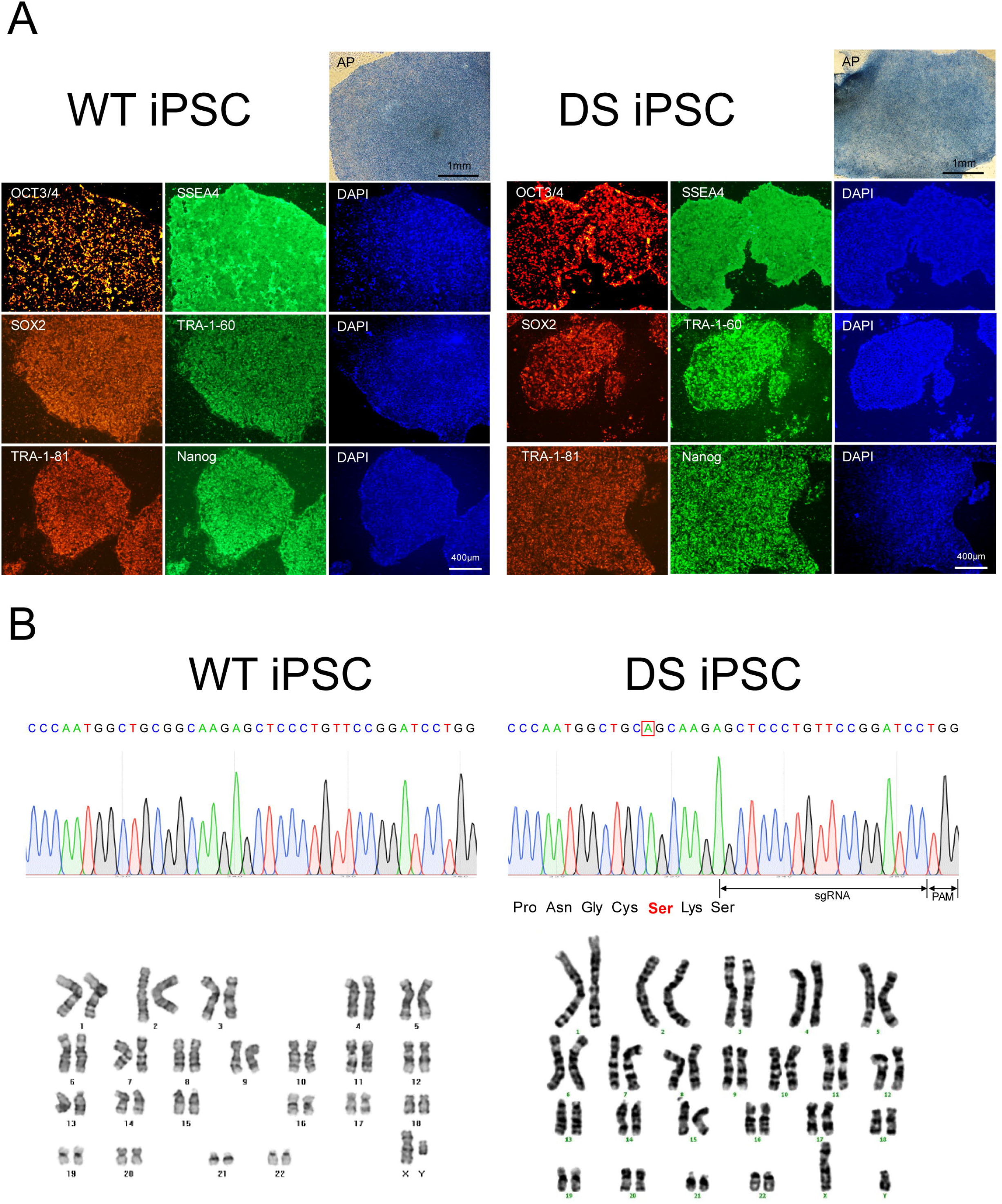
Specific marker and genotypical characterization of fibroblast-derived iPSC. Control (WT) and fibroblast from ALD patient (DS) were transduced with sendai viral vectors expressing reprogramming factors Oct, Sox2, Klf4, and c-Myc **A** Alkaline phosphatase staining and immunofluorescence analysis, including OCT3/4, SOX2, TRA-1-81, SSEA4, TRA-1-60 and NANOG markers, shows the pluripotency of WT and DS iPSC. **B** Sanger sequencing of WT and DS iPSC confirmed c.1534G>A (p.Gly512Ser). iPSCs, originated from WT and DS fibroblast, presented normal chromosomal number and structure.

### Efficient gene correction via ssODN-mediated HDR

We introduced the CRISPR/Cas9 system with an 184bp ssODN donor to DS iPSCs (Fig 2A). The double-strand break point upon introducing the Cas9 protein was located 22 bp upstream of the mutation site. Fig 2B shows a schematic illustration of gene correction in iPSCs. DS iPSCs were co-transfected with SpCas9 (2 μg), pRGEN_ABCD1 (4 μg), and ssODN (500 pmol). After 3 h of puromycin selection, 100 to 150 colonies survived; these colonies were cultured for 10 days, then triturated and transferred separately to 24-well plates. Triturated colonies in 24-well plates were cultured for 4 more days. Then, half of each colony in one well was used for gDNA extraction and PCR amplification (amplicon size: 742bp). We used *Apa1* digestion for positive selection in cultured colonies. Positive colonies were confirmed by Sanger sequencing. Sequencing results showed a mixed population of corrected and uncorrected cells. In each positive well, colonies were separated once more into 24-well plates, cultured for 4 days, and half of each colony in a single well was collected for Apa1 digestion and Sanger sequencing. Finally, we obtained 3 isogenic gene-corrected iPSC colonies (DSC1, DSC2, DSC3) after just two subculture passages (Fig 3A). We performed immunocytochemistry assay and karyotyping of gene-corrected iPSCs. They expressed pluripotency markers and normal karyotypes without abnormalities in the number or structure of chromosomes (Fig 4).

**Fig 2.**
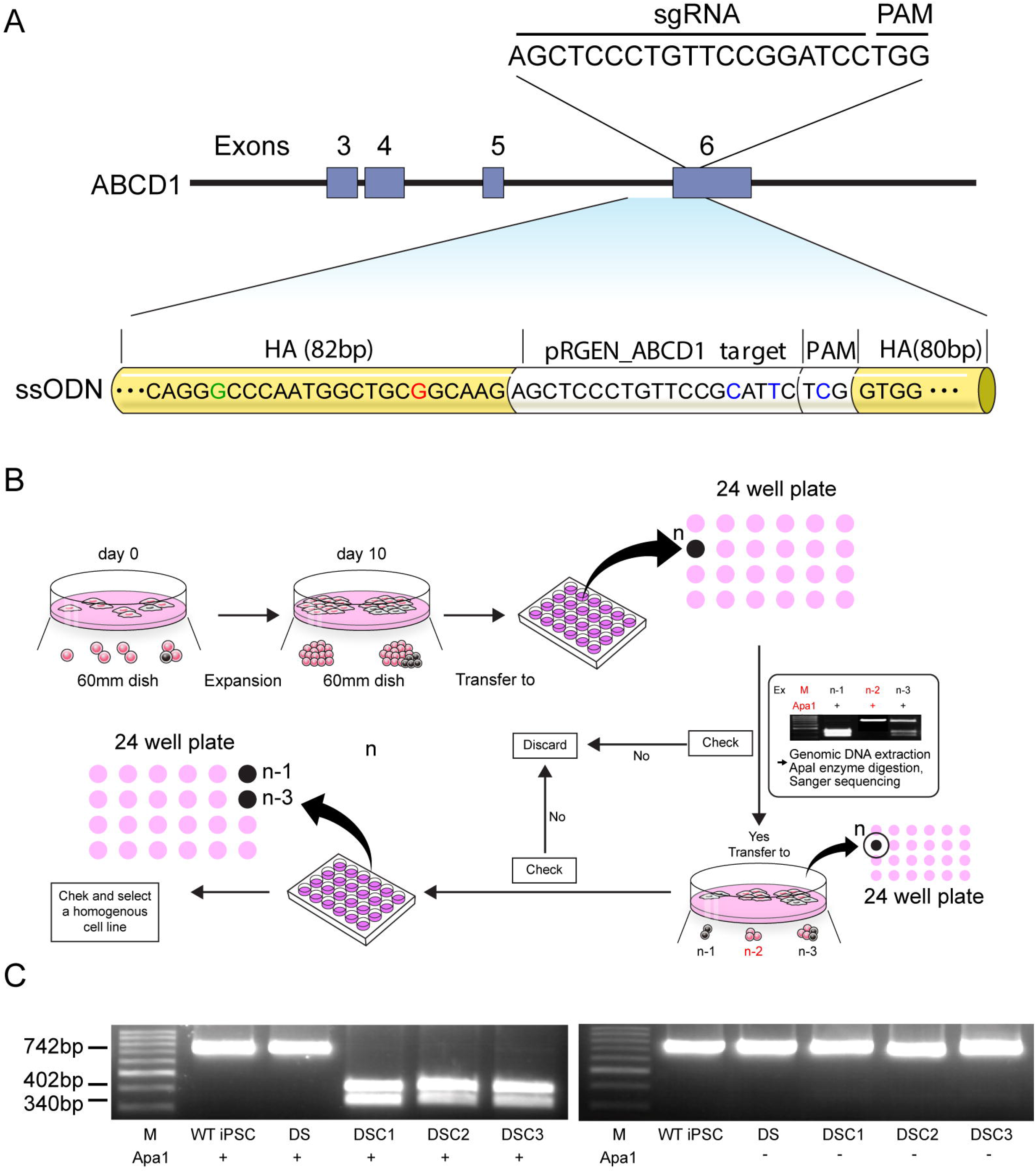
Gene correction of patient-derived iPSCs. **A** Schematic diagram of gene correction with the Cas9 system and single-stranded oligodeoxynucleotides (ssODN). The sgRNA was targeted near the mutation site (c.1534G>A), and the ssODN sequence had five base pair differences from the patient’s genomic sequence, included one for Apa1 enzyme digestion, one for gene correction (A>G), and three for preventing re-cleavage by the Cas9 protein. **B** Schematic illustration of the serial passage method for obtaining isogenic mutation-corrected iPSCs. **C** Amplified PCR products (amplicon size: 742 bp) of subcultured colonies were digested by the *Apa1* restriction enzyme. If the mutation site has been corrected, the PCR amplicon would be digested into 402-bp and 340-bp segments by the *Apa1* restriction enzyme. Three gene-corrected iPSC colonies (DSC1, DSC2 and DSC3) showed positive results without the undigested band.

**Fig 3.**
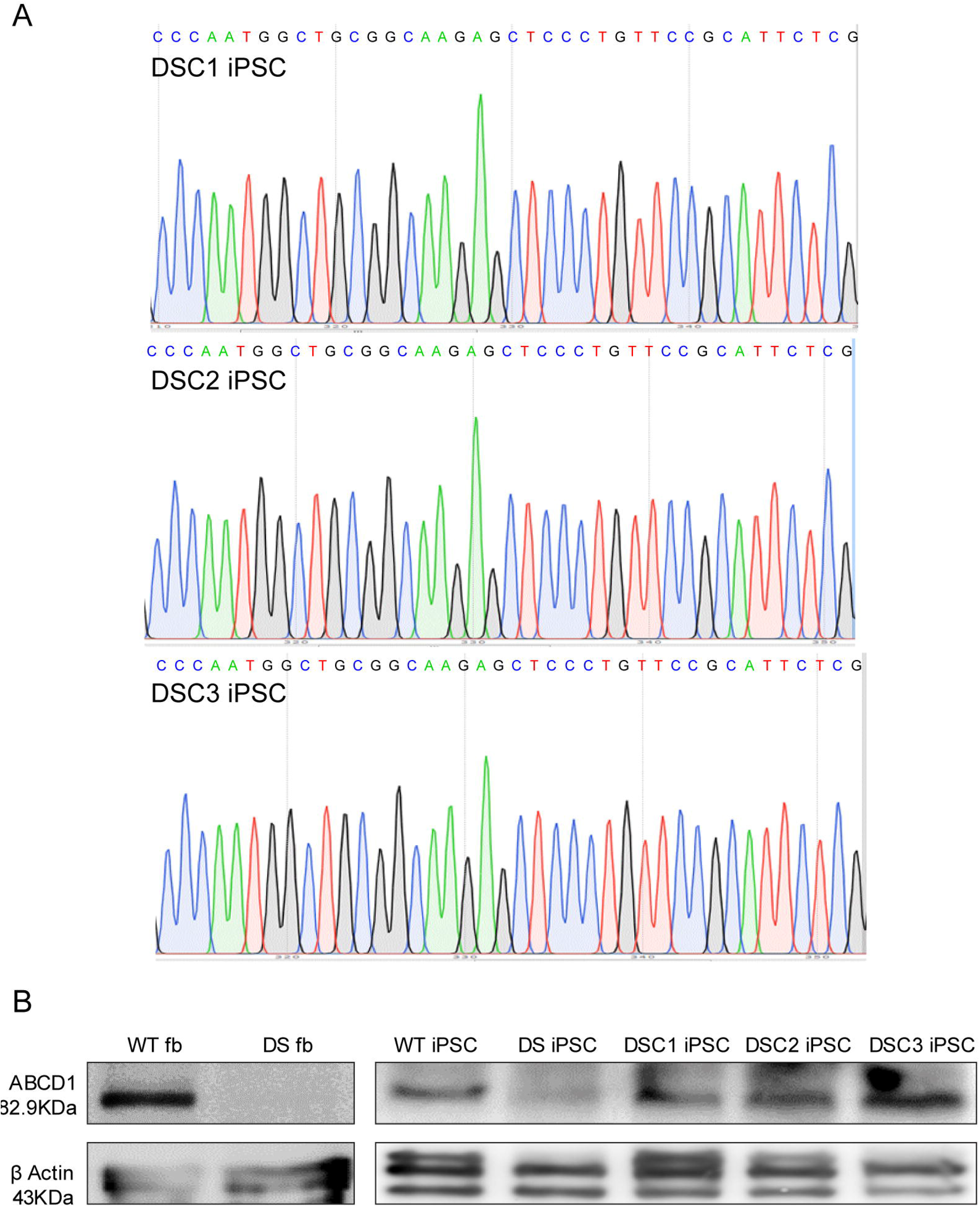
Correction of mutation site in patient-derived iPSC rescue ALDP expression. **A** Sanger sequencing confirmed that the mutation site (c.1534G>A, gray bar) had been corrected. Red squares represent artificial silent mutation sites. **B** Western blotting detected ALDP expression in the DSC1, DSC2, and DSC3 cell lines after gene correction. No ALDP was detected in ALD patient (DS) fibroblasts and DS iPSCs. Normal fibroblasts (WT) and WT iPSCs were used as positive controls.

**Fig 4.**
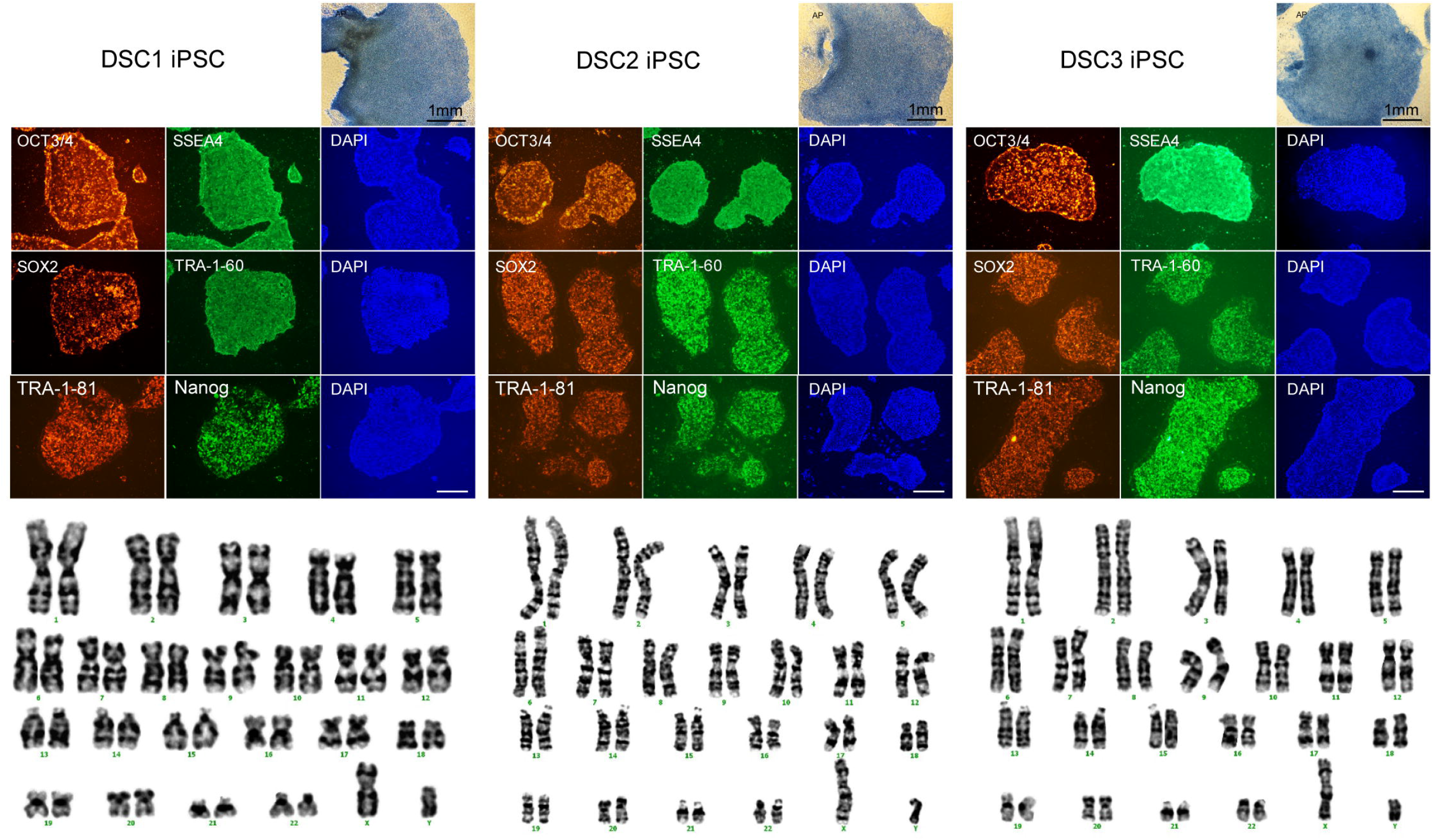
Gene-corrected iPSC lines expressed pleuripotency and normal karyotypes. Three gene-corrected iPSC colonies (DSC1, DSC2 and DSC3) expressed positive of alkaline phosphatase (AP) staining and immunocytochemistry stain for pleuripotency markers, including OCT3/4, SOX2, SSEA4, TRA-1-60 and NANOG markers. Karyotypes presented normal chromosomal number and structure.

### ALDP expression after gene correction

We performed western blotting to validate *ABCD1* expression after gene correction (Fig 3B). No ALD protein (ALDP) expression was observed in DS fibroblasts and DS iPSCs. ALDP expression was confirmed in the DSC1, DSC2, and DSC3 cell line, which were the mutation-corrected iPSC lines. The DSC1 cell line was maintained for 33 passages, DSC2 for 23 passages, and DSC3 for 8 passages.

### Off-target analysis

Cas-OFFinder captured one on-target site and one 2-bp mismatch off-target site; no 1-bp mismatch off-target site was found without a DNA bulge. If DNA bulges were permitted, there were 24 1- or 2-bp mismatch off-target sites (S1 Table). We performed WES to investigate the overall off-target effects of CRISPR/Cas9 introduction. The number of single nucleotide polymorphisms (SNPs) and variants from gene-corrected iPSC cell lines were similar to DS iPSCs (Table 1). We analyzed the on-target reads of DSC1, DSC2, and DSC3 iPSCs in comparison with untreated iPSCs (DS iPSCs). Eight significant variants were captured for DSC1, two for DSC2, and none for DSC3 (S1 Table). These variants were not in crucial genes (S2 Table). Furthermore, the pathogenic evidence categories of these variants were moderate or lower, based on the American College of Medical Genetics and Genomics (AGMG) guidelines.

**Table 1.**
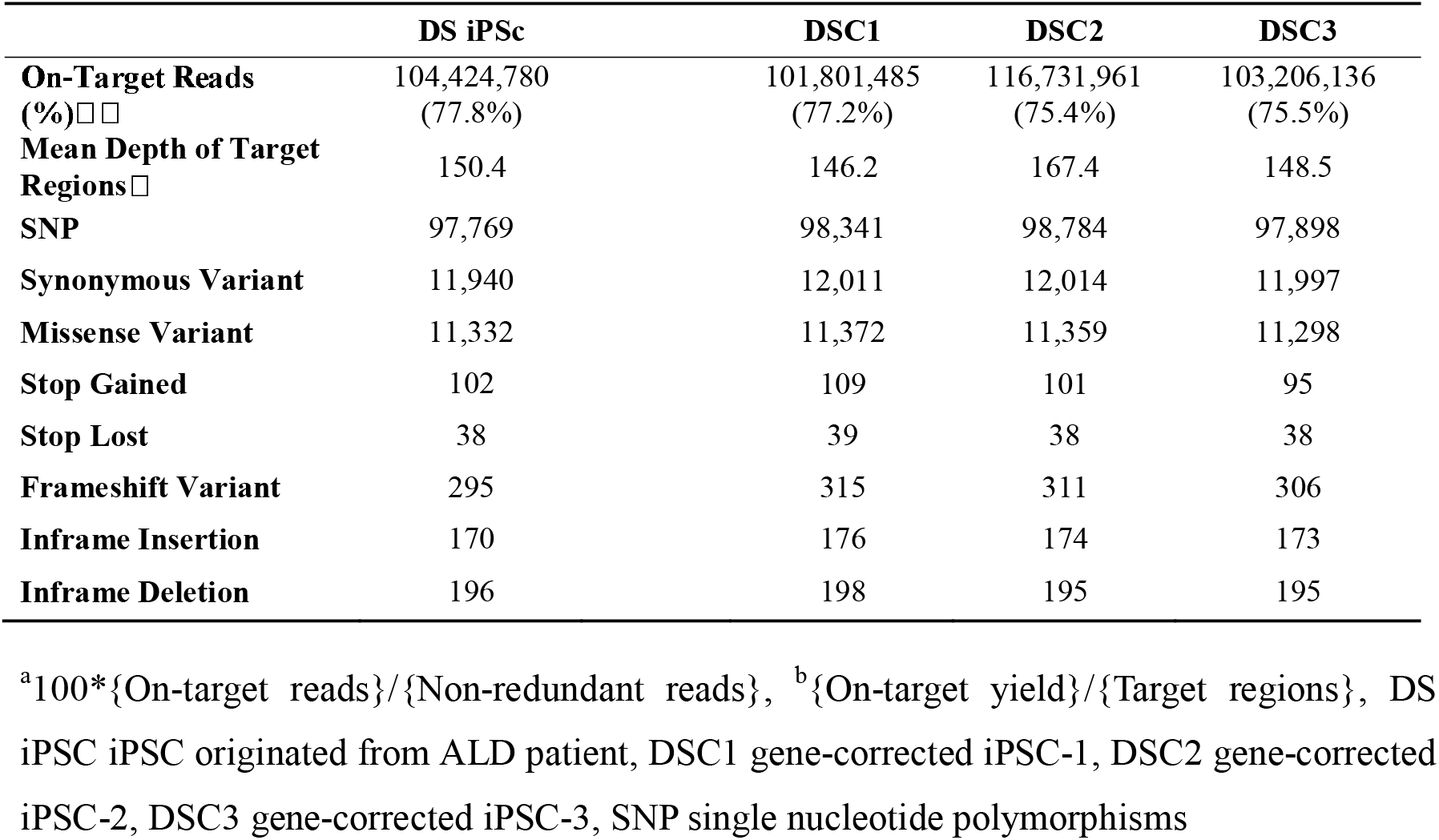
The whole exome sequencing (WES) results of control and mutation corrected iPSC cell lines

### Differentiation of oligodendrocytes from mutation-corrected ALD patient iPSCs

For generation of neural precursor cells (NPCs), 2.8 x 10^5^ cells of mutation-corrected iPSCs were dissociated to single cell and seeded on Matrigel coated 6 well plate. These were cultured with neuronal induced medium within DMEM/F12, 1x non-essential amino acids, 1x N2 and 1x B27 supplements, 10μM SB, 1 μM LDN193189, 25μg/ml human insulin and 100nM RA. After 8 days, the medium was changed to neuronal induced medium with 1μM SAG for patterning to gliaform NPCs.

To generate the O4 positive intermedia OLs, cultured cells were dissociated and plated on new plates. Then, cells were transduced with mSOX10 lentivirus, then medium was changed to ODM, that is NIM with 1 mg/mL of doxycycline, 10 ng/ml of IGF1, 5ng/ml of HGF, 1μM of cAMP, 10ng/ml of NT-3, 60ng/ml of T3, 10ng of PDGFaa and 100ng/ml of biotin. Transduced cells were maintained for 10-12 days. Thereafter cells expressed O4 and MBP (Fig 5).

**Fig 5.**
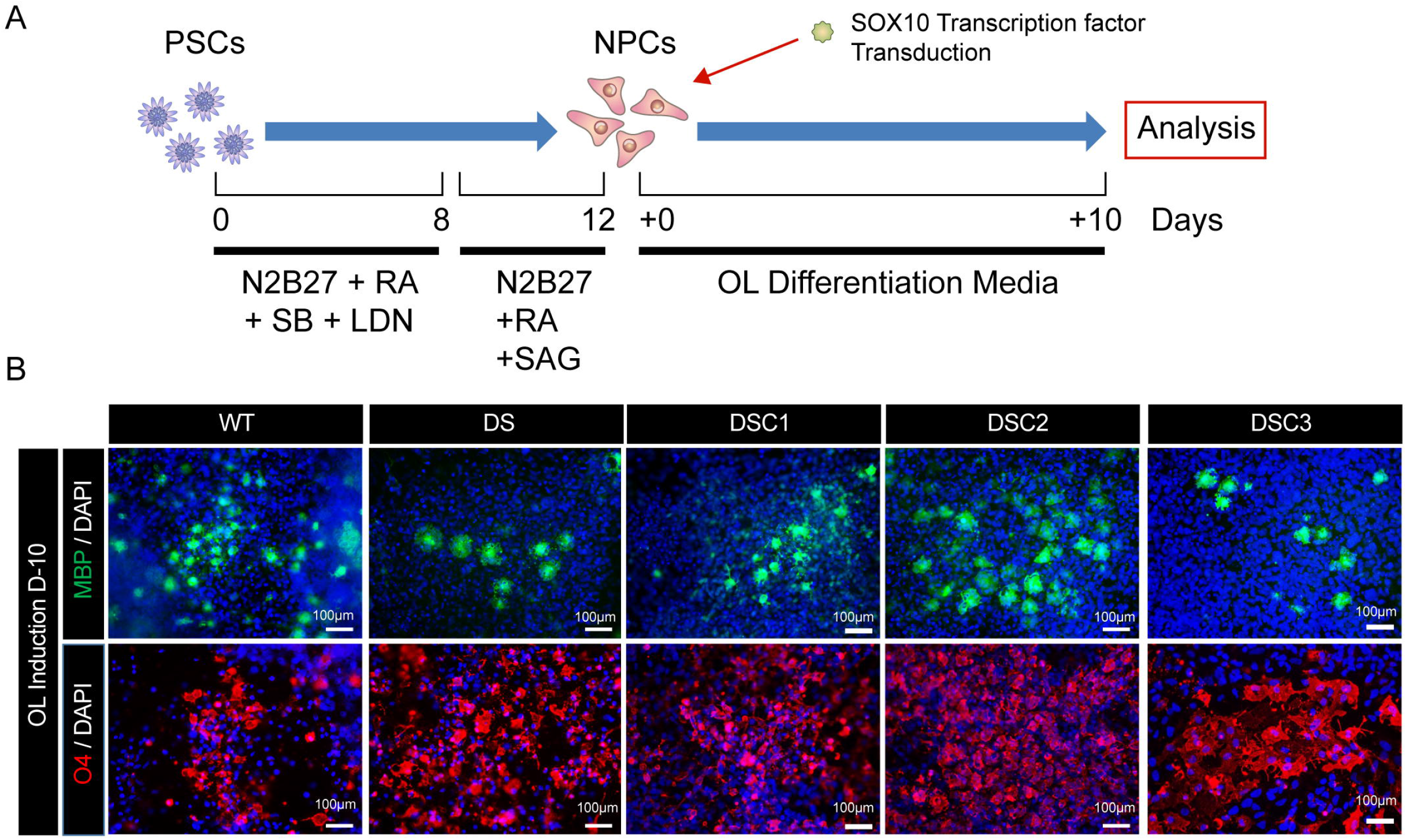
Differentiation of intermediate oligodendrocytes from iPSc. **A** Scheme of oligodendrocyte (OL) differentiation with lentiviral transduction B Successful differentiation of intermediate oligodendrocytes from control (WT), ALD patient (DS) and mutation-corrected (DSC1, DSC2 and DSC3) iPSCs. There was double positive staining of MBP and O4.

### Analysis of saturated very long chain fatty acids and LPCs C26:0

After intermediate OLs differentiation, we verified the improvement in VLCFA metabolism. Previously we reported that VLCFA level was low in ALD patient-derived iPSCs and metabolic derangement of VLCFA was discovered after oligodendrocyte differentiation [8]. We MACs sorted O4 positive cells 22 days after transduction by using Anti-O4 MicroBeads and LS columns. 2 x 10^5^ cells per each sample were analyzed for VLCFAs and LPC VLCFAs.

VLCFAs (C22:0, C24:0, C26:0) were no significant differences between wild, DS and DSC OLs. VLCFA metabolic ratios, too (Kruskal-Wallis test; C24:0/C22:0, p=0.552; C26:0/C22:0, p=0.687). C24:0 LPC and C26:0 LPC decreased comparing as DS OLs, but it wasn’t statistically different (Kruskal-Wallis test; C24:0 LPC, p=0.072; C26:0 LPC, p=0.149).

## Discussion

ALD is an inherited monogenic metabolic disease, but there is no curative therapeutic option. Until now, attempts to overcome this genetic disease have been to reduce VLCFA through an alternative pathway such as ABCD2 or ABCD3, or to control oxidative stress or inflammatory processes, the output of the gene mutation [13–15]. Although these strategies are effective *in vitro* or *in vivo,* they fail to prove effective at the clinical level. This study demonstrates the first ALD patient-derived iPSC lines in which the disease-causing mutation has been corrected, and these cell lines exhibit normal ALDP expression.

With the introduction of the CRISPR/Cas9 system, the efficiency of genome editing has increased remarkably. It has become possible to delete a gene fragment, correct a mutation site, insert a large gene fragment, and enhance or repress gene expression through the CRISPR/Cas9 System [9]. This technology can reveal the pathophysiology of an unknown disease or overturn previous theories. Genome editing using the CRISPR/Cas system can correct genetic mutations via non-homology-mediated end joining or HDR [10]. Genome editing via HDR can correct a mutation site, but this depends on the cell cycle, and HDR efficiency is very low, particularly in neuronal cells [16]. To overcome this efficiency problem, a homology-independent targeted insertion was developed; however, with this method, the indel can remain at the integration site and *in vivo* efficiency is low [17]. There are two strategies to obtain genome-edited iPSCs; genome editing in cells before inducing them to become iPSCs, or in iPSC directly. We first used human fibroblast for gene correction. Preferred cleavage targets for HDR were selected, and cleavage efficiency was measured in 293T and human fibroblast cells. However, there was a difference in cleavage efficiency between 293T and human fibroblast (data not shown). Generally, the result from human fibroblasts was lower than that seen in 293T cells. Substitution via HDR mechanism in fibroblast is too difficult due to low HDR efficiency. Furthermore, for the generation of gene-corrected iPSC lines, the selected cells should be treated with Yamanaka factors. During this step, many cells may die, and there is the risk of undesirable modifications being incorporated. The efficiency of genome modification using iPSCs is extremely low and the experimental procedure is labor-intensive [18]. In this experiment, only 2% (3/150) colonies were positive for Apa1 enzyme digestion initially. Because the HDR rate is extremely low, single cell culture for obtaining isogenic colonies is expensive and labor-intensive. Therefore, we used the serial passage method. Following this method, we succeeded in obtaining isogenic mutation-corrected cell lines after just two subcultures. All steps were performed within a duration of 1 month.

Although ALD is a genetic disease, there is no genotype-phenotype correlation. Increased VLCFA levels due to the *ABCD1* mutation has no correlation with phenotype and disease severity. It is possible that other factors, such as epigenetic or environmental factors, affect disease manifestation [19]. Blood brain barrier fragility, metabolic derangement, and a malfunctioning immune system are also potential secondary factors [20–22]. In spite of this ambiguity, all ALD patients possess the *ABCD1* mutation. Although ALDP is a homodimeric protein, *ex vivo* delivery of *ABCD1* cDNA is successful at the research and clinical levels [23, 24]. *In vivo* delivery of *ABCD1* cDNA via adeno-associated virus (AAV) has been performed, and also showed positive results [25]. However, *ex vivo* gene therapy has a limited treatment indication; it must be performed in the early phase of cerebral inflammation. *In vivo* gene therapy has the potential risks of viral toxicity and gene fragment integration, and its el?icacv is temporary. In this study, we efficiently generated isogenic mutation-corrected iPSC lines, all of which exhibited normal ALDP expression. Off-target analysis indicates that these strategies would be safe, without genotoxicity due to the introduction of Cas9 and ssODN. In particular, the DSC3 iPSC line showed no significant differences compared to the DS iPSC line. *In silico* off-target analysis cannot reflect real world experimental results. In this experiment, WES data from post gene-corrected iPSC lines aligned with those from the untreated iPSC cell line. This method is better than unbiased off-target analysis. Variants with significant number were identified and compared with control to evaluate clinical significance.

We first expected that newly expressed ALDP would work in oligodendrocytes. Previously we couldn’t find significant differences between OPCs from ALD iPSc, AMN iPSc and healthy[8]. Therefore, we differentiated mutation-corrected iPSc to oligodendrocytes. We successfully obtained O4 positive oligodendrocytes, then analyzed VLCFA and LPC VLCFA profiles. There was no significant difference in VLCFA, but LPC showed decreased amount of LPC C24:0 and LPC C26:0. However, it wasn’t statistically significant. We hypothesized two reasons. At first, we analyzed O4 positive oligodendrocytes. In developing brain, fatty acid enzyme activity is rapidly increased at the active myelinating phase [26]. ALDP works as consuming the excessive VLCFA. In oligodendrocytes with O4 positive and MBP positive staining, myelination starts. Maybe the VLCFA differences between normal and ALD would become clear after active myelination phase. Second, we analyzed small numbers of each samples. There is a limit of massive oligodendrocyte differentiation in laboratory level.

In conclusion, we first generated ALD patient-derived iPSCs, and subsequently corrected the disease-causing mutation. This rescued normal ALDP expression in mutation-corrected iPSC lines. However, it did not result in the improvement of abnormal VLCFA metabolism. In the future, the mutation corrected iPSCs will be used for research on ALD pathophysiology and for the development of therapeutic options.

## Acknowledgements

We thank MID (Medical Illustration & Design), a part of the Medical Research Support Services of Yonsei University College of Medicine, for all artistic support related to this work.

