## Supplementary Tables for "Successful Correction of ALD Patient-derived iPSCs Using CRISPR/Cas9"

**Supplementary materials**

**Supplementary Sequences**

Sequences of single-stranded oligonucleotide donor templates (ssODN) used in HDR: TCTCTGGCGTCAGCGGCTGTTGCCCCTGCAGGTGGAGGAAGGCATGCATCTGCTCATCACAGGGCCCAATGGCTGCGGCAAGAGCTCCCTGTTCCGCATTCTCGGTGGGCTCTGGCCCACGTACGGTGGTGTGCTCTACAAGCCCCCACCCCAGCGCATGTTCTACATCCCGCAGAGGTAAGGA

Red: sgRNA sequence

Gray: silent mutation for ApaI restriction enzyme

Yellow: to prevent re-cleavage by sgRNA-Cas9 complex

Underline: PAM site

**Supplementary Tables**

Supplementary table 1. *In silico* analysis of off-targets in human genome [GRCh38] (http://www.rgenome.net/cas-offinder/)

| **Bulge type** | **Target of genomic DNA** | **Chromosome** | **Position** | **Mismatches** |
| --- | --- | --- | --- | --- |
| No | AGCTCCCTGTaCCGGATtCAGG | chr12 | 94351279 | 2 |
| No | AGCTCCCTGTTCCGGATCCTGG | chrX | 153740142 | On target |
| DNA | AGCTCCCTGTaCCGGATTCaGGG | chr12 | 94351278 | 2 |
| DNA | AGCTCCCTGTaCCGGATTCaGGG | chr12 | 94351278 | 2 |
| DNA | AGCTCaCTGTTCtGGCATCCTGG | chr12 | 48050269 | 2 |
| DNA | AGGCTCCCTGTTCCcGATgCTGG | chr7 | 138756754 | 2 |
| DNA | AGGCTCCCTGTTCCcGATgCTGG | chr7 | 138756754 | 2 |
| DNA | AGCTCtCTGTTCCAGGATgCAGG | chr5 | 179452005 | 2 |
| DNA | AGCTCCCGgGTTCCGGcTCCTGG | chr16 | 50859724 | 2 |
| DNA | AGCTCCCgGGTTCCGGcTCCTGG | chr16 | 50859724 | 2 |
| DNA | AGCTCCCgGGTTCCGGcTCCTGG | chr16 | 50859724 | 2 |
| DNA | gGCTCCCTGATTCaGGATCCAGG | chr21 | 35632569 | 2 |
| DNA | AGaTGCCCTGTTCCGGATtCAGG | chr10 | 70145162 | 2 |
| DNA | ATaCTCCCTGTTCCaGATCCAGG | chr10 | 19476466 | 2 |
| DNA | AGCTCTCCTGcTCCGaATCCAGG | chr10 | 116053719 | 2 |
| DNA | AtACTCCCTGTTCCaGATCCAGG | chr10 | 19476466 | 2 |
| DNA | AGCTCCCTGTTCCGGcTgGCAGG | chr6 | 169918304 | 2 |
| DNA | AGCCTCCCTGTTCCaGAaCCTGG | chr6 | 41000289 | 2 |
| DNA | AGCTCCCTtTTCCtGATACCAGG | chr6 | 57243222 | 2 |
| DNA | AGCTCCCTGTTCCGGcTGgCAGG | chr6 | 169918304 | 2 |
| DNA | AGCCTCCCTGTTCCaGAaCCTGG | chr6 | 41000289 | 2 |
| DNA | AGCTCCCTGTTCCGGATCCtGGG | chrX | 153740142 | 1 |
| DNA | AGCTCCCTGTTCCGGATCCtGGG | chrX | 153740142 | 1 |
| DNA | AGCTCCCTGTTCCGGATcCtGGG | chrX | 153740142 | 2 |
| DNA | gAGCTCCCTGTTCCGGATCCTGG | chrX | 153740141 | 1 |
| DNA | gaGCTCCCTGTTCCGGATCCTGG | chrX | 153740141 | 2 |

Red: mismatched base pairs, compared with sgRNA sequences (AGCTCCCTGTTCCGCATTC)

Supplementary table 2. Captured variants of DSC1, DSC2 and DSC3, compared to untreated (DS) iPSc in WES

| **Chromosome: position** | **Gene** | **HGVSc** | **HGVSp** | **Frequency** | **Depth** | **ACMG** |
| --- | --- | --- | --- | --- | --- | --- |
| **DSC1** |  |  |  |  |  |  |
| chr22:26902308-26902308 | *TFIP11* | c.481G>T | p.Gly161Cys | 0.462 | 91 | PM2,PP3 |
| chr1:22839448-22839448 | *ZBTB40* | c.2493C>T | p.Phe831= | 0.499 | 387 | PM2 |
| chr15:101983785-101983785 | *PCSK6* | c.376C>A | p.His126Asn | 0.329 | 76 | PM2 |
| chr19:24115169-24115169 | *ZNF726* | c.251A>G | p.Asp84Gly | 0.613 | 31 | PM2 |
| chr2:202568864-202568864 | *ALS2* | c.4916T>G | p.Ile1639Arg | 0.178 | 45 | PM2 |
| chr2:202568866-202568866 | *ALS2* | c.4914T>G | p.Gly1638= | 0.186 | 43 | PM2 |
| chr3:141678649-141678649 | *TFDP2* | c.738T>G | p.Phe246Leu | 0.244 | 86 | PM2 |
| chr4:2337476-2337476 | *ZFYVE28* | c.447C>T | p.Tyr149= | 0.498 | 229 | PM2 |
| **DSC2** |  |  |  |  |  |  |
| chr13:47470831-47470831 | *HTR2A* | c.138A>C | p.Ser46= | 0.2 | 85 | PM2 |
| chr17:19687101-19687101 | *ULK2* | c.2369T>G | p.Leu790Arg | 0.5 | 48 | PM2 |

HGVSc, the Human Genome variation Society coding sequence name; HGVSp, the Human Genome variation Society protein sequence name; ACMG, the American College of Medical Genetics and Genomics
